# Profiling serum immunodominance following SARS-CoV-2 primary and breakthrough infection reveals distinct variant-specific epitope usage and immune imprinting

**DOI:** 10.1101/2024.08.14.607882

**Authors:** Jeffrey Seow, George C.E. Jefferson, Michael Keegan, Yeuk Yau, Luke B. Snell, Katie J. Doores

## Abstract

Over the course of the COVID-19 pandemic, variants have emerged with increased mutations and immune evasive capabilities. This has led to breakthrough infections (BTI) in vaccinated individuals, with a large proportion of the neutralizing antibody response targeting the receptor binding domain (RBD) of the SARS-CoV-2 Spike glycoprotein. Immune imprinting, where prior exposure of an antigen to the immune system can influence the response to subsequent exposures, and its role in a population with heterogenous exposure histories has important implications in future vaccine design. Here, we develop an accessible approach to map epitope immunodominance of the neutralizing antibody response in sera. By using a panel of mutant Spike in a pseudovirus neutralization assay, we observed distinct epitope usage in convalescent donors infected during wave 1, or infected with the Delta, or BA.1 variants, highlighting the antigenic diversity of the variant Spikes. Analysis of longitudinal serum samples taken spanning 3 doses of vaccine and subsequent breakthrough infection (BTI), showed the influence of immune imprinting from the ancestral-based vaccine, where reactivation of existing B cells elicited by the vaccine resulted in the enrichment of the pre-existing epitope immunodominance. However, subtle shifts in epitope usage in sera were observed following BTI by Omicron sub-lineage variants. Antigenic distance of Spike, time after last exposure, and number of vaccine boosters may play a role in the persistence of imprinting from the vaccine. This study provides insight into RBD neutralizing epitope usage in individuals with varying exposure histories and has implications for design of future SARS-CoV-2 vaccines.

**Author Summary:** Throughout the COVID-19 pandemic, the continued emergence of new SARS-CoV-2 variants has resulted in a rise in breakthrough infections (BTIs). Infection with different variants has led to varying exposure histories in the general population. Although the neutralizing response to Spike has been thoroughly characterized, with several key epitopes identified, there is a lack of knowledge of the proportion each epitope contributes to the neutralizing response in sera and how this is affected by exposure history. Here, we use a panel of mutant Spike pseudoviruses to screen epitope usage and immunodominance in polyclonal sera. In a cohort of unvaccinated donors infected with different variants, distinct epitope usage was observed, highlighting the antigenic diversity between the variant Spikes. Furthermore, samples collected spanning multiple vaccine doses and BTI showed the influence of prior immunity from the vaccine on epitope usage. Although a large proportion of the immune response following BTI could be attributed to enrichment of pre-existing immunodominance from the vaccine, subtle shifts in epitope usage were observed with infection by more mutated variants. This work gives more detailed insight into differences in the neutralizing response of individuals with varying exposure histories that may inform next generation SARS-CoV-2 vaccines.

## Introduction

The SARS-CoV-2 Spike is the major target of neutralizing antibodies elicited following infection and vaccination[1]. Several distinct immunodominant epitopes have been identified on the receptor binding domain (RBD)[2–7], and to a lesser extent on the N-terminal domain (NTD) [2,8,9]. The selective pressure on the virus from the humoral immune response has driven the emergence of new variants over the course of the COVID-19 pandemic. These variants have evolved multiple mutations in Spike resulting in increased viral immune escape and enhanced transmission. For example, the recent XBB.1.5 and BA.2.75 variants have been shown to have greatly reduced neutralization by COVID-19 vaccinee sera[10,11]. Although neutralizing epitopes on Spike have been well characterised through structural analysis, the contribution of antibodies targeting these defined epitopes make in polyclonal sera on overall neutralizing activity have not been extensively studied[12]. The emergence of new SARS-CoV-2 variants, able to successfully evade antibody responses, has led to an increase in breakthrough infections (BTI) and a wide heterogeneity of SARS- CoV-2 exposure histories[13]. Determining the predominant neutralizing epitopes targeted on Spike (i.e. the epitope immunodominance) following different vaccination and infection histories is important for understanding the immune selective pressures on Spike and potential mutations that may arise in future SARS-CoV-2 variants of concern, as well as providing insights into potential population susceptibility to newly arising variants.

Immune imprinting is the phenomenon in which the prior exposure of an antigen to the immune system can influence the response to subsequent exposures to related antigens. Several studies have indicated that immune imprinting limits *de novo* antibody responses against an infecting SARS-CoV-2 variant in vaccinated donors. For example, recent studies have observed a high frequency of cross- reactive B cells and a low frequency of variant specific B cells in PBMC of individuals experiencing BTI suggesting that *de novo* responses specific for the infecting variant are suppressed[14–18]. Furthermore, antigenically distant variant booster vaccinations have been shown to reactivate existing B cell responses from WT vaccination, preventing the generation of variant-specific mAbs, potentially impeding immunity to emerging variants[17,19–22]. These observations are supported by an increase in antibody somatic hypermutation in donors experiencing BTI, implying continued affinity maturation of existing B cells[3,14,22,23]. Although these studies confirm the influence of immune imprinting from Wuhan-1-based vaccines on subsequent variant boosting or infection, further characterisation of this response is lacking in larger cohorts. Furthermore, how immune imprinting might be impacted by the SARS-CoV-2 variant at the time of first exposure has not been considered in great detail[24].

Several techniques have been employed to map the Spike epitopes targeted by neutralizing antibodies generated following infection and vaccination. Traditional epitope mapping by peptide array, although high-throughput, can only identify linear epitopes and therefore may miss quaternary epitopes[25]. More recent approaches include mAb isolation[2,3,23], electron microscopy polyclonal epitope mapping (EMPEM)[26], 10x genomics[14,22,27], and deep mutational scanning (DMS)[28–31]. Although these approaches provide epitope mapping at high resolution, they are limited by the cohort size that can be examined, and some require access to peripheral blood mononuclear cells (PBMCs). Here, we have developed a more accessible approach to serum epitope mapping utilizing an engineered neutralization resistant Spike based on XBB.1.5 and a panel of mutated Spikes where selected epitopes were reverted back to the ancestral (wildtype, WT) sequences. Restoration of neutralization allowed identification and comparison of immunodominant epitopes targeted in sera from a primary infection cohort (including wave 1, Delta and BA.1 infections) and a longitudinal vaccination cohort (including three doses of BNT162b2 mRNA vaccine) where individuals also subsequently experienced a BTI. Using this approach, we show that primary infection with variants elicit distinct immunodominance profiles compared to a Wuhan-1 based vaccine. Additionally, the antibody response to BTI is imprinted by prior vaccination, resulting in maturation of pre-existing epitope usage. As the virus continues to evolve, understanding shifts in the immunodominant epitopes on Spike are crucial for predicting future variants and aid in the design of optimized vaccines.

## Results

### Design of highly immune evasive mutant XBB.1.5+Spike and virus revertant panel

Introduction of Spike mutations in the epitopes of known SARS-CoV-2 nAbs to WT Spike in pseudotyped neutralization assays has been used to identify epitopes of mAbs and map neutralization specificity in sera[20,32–34]. However, the polyclonal nature of the nAb response to SARS-CoV-2 can dampen the effect of individual point mutations on neutralization reduction. We therefore took an alternative approach where we used a highly neutralization resistant Spike as a template and generated a panel of virus Spike revertants where selected epitopes were mutated back to the ancestral strain. Rescue of serum neutralization against a revertant virus would indicate presence of neutralizing antibodies against this epitope that could be quantified.

In early 2023, XBB.1.5 was the dominant strain of SARS-CoV-2 in the UK. The Spike of XBB.1.5 differed from the ancestral Wuhan-1 Spike by 40 amino acid mutations, resulting in a large reduction (29-fold) in neutralization potency for fourth-dose monovalent mRNA vaccine sera[10,35,36]. Sequence alignment of other highly mutated endemic SARS-CoV-2 variants identified near to that time, BQ.1.1 and BA.2.75, showed similar Spike mutations to XBB.1.5 (**Figure. 1A**) but these variants also encoded additional RBD mutations. To generate a more immune evasive strain, the K444T and L452R mutations present in BQ.1.1 and BA.2.75 were incorporated into the XBB.1.5 sequence and the Spike variant was named “XBB.1.5+”. To determine the neutralization resistance of XBB.1.5+, a small panel of vaccinee (3 doses of BNT162b2 vaccine) and BTI sera were tested against D614G, XBB.1.5 and XBB.1.5+. A 25-fold decrease in GMT ID_50_ for XBB.1.5 compared to D614G was observed and further 2-fold decrease was observed for XBB.1.5+ compared to XBB.1.5 (**Figure 1B**).

**Figure 1:**
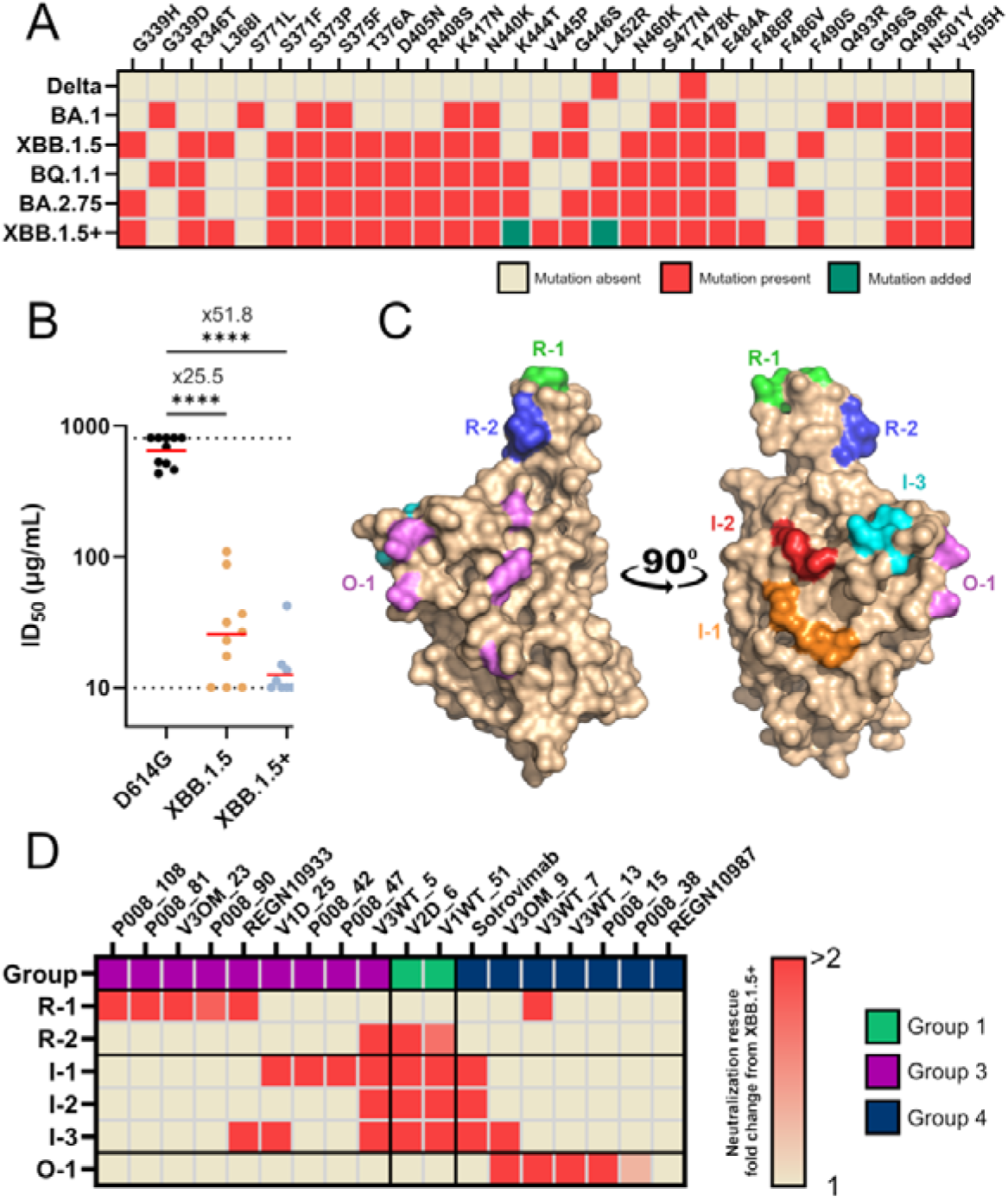
Generation and validation of XBB.1.5+ mutant and virus revertant panel. (A) Amino acid mutations present in the RBD of immune evasive SARS-CoV-2 variants Delta, BA.1, XBB.1.5, BQ.1.1 and BA.2.75, and RBD mutations present in engineered XBB.1.5+. (B) Comparison of serum neutralization ID_50_s against D614G, XBB.1.5 and XBB.1.5+ pseudotyped virus. Sera are collected from individuals who had had 3 doses of BNT162b2 mRNA vaccine and a BTI (Kruskal-Wallis multiple comparison test). (C) Surface representation of the WT Spike RBD (pdb: 6XM0) with amino acids reverted back to WT indicated by coloured surface. Mutations shared in one revertant are coloured the same. A full list of mutations can be found in Supplementary Table 2. (D) Heatmap showing fold rescue of neutralisation for reverted viruses compared to XBB.1.5+ for a panel SARS-CoV-2 mAbs isolated from convalescent and/or vaccinated donors[2,3,23] as well as therapeutic mAbs Sotrovimab, REGN10933 and REGN10987. mAbs were previously clustered using a competition binding ELISA into three RBD-specific competition groups[2,3,23].

Having identified a highly resistant Spike, primers were designed to mutate individual residues in the XBB.1.5+ template back to the amino acids present in the ancestral sequence (**Supplementary Table 1**). As the majority of the neutralizing activity in sera is directed against the RBD[37,38], the Spike revertants focused on mutations in RBD only. Overall, six mutated Spikes were generated by grouping together mutations in close proximity to widen the range of potential neutralization rescue whilst retaining resolution in epitope mapping (**Figure 1C, Supplementary Table 2** ). Epitopes R-1 and R-2 were in vicinity of the receptor binding motif (RBM) which binds to ACE2. I-1, I-2 and I-3 include epitopes on the interior face of the RBD, which are hidden when the RBD is positioned in the “down” conformation. The last epitope, O-1, was located on the outer face of RBD and is accessible when RBD is in both the “up” and “down” conformations.

### Neutralization rescue of mAbs against virus revertant panel

To verify whether antibody neutralisation can be rescued in the XBB.1.5+ virus revertant panel, and to gain insight into the types of serum antibodies that facilitate neutralization, monoclonal antibodies (mAbs) isolated and characterised from infected[2] and BTI[23] donors were tested (**Figure 1D** ). This panel of mAbs have previously been categorised into epitope groups based upon competition binding assays[2,39,40]. Therapeutic mAbs Sotrovimab and Ronapreve (a cocktail containing REGN10987/Imdevimab and REGN10933/Casirivimab), whose epitopes have been defined using structural biology, were also tested. Neutralization rescue was observed for all mAbs for at least one revertant Spike. Group 3 mAbs, which have been previously shown to bind to the RBM and compete with ACE2 binding of Spike, had neutralization rescue with epitope revertant R1. This included therapeutic mAb REGN10933 for which rescue by R-1 and I-3 epitope reversions are supported by previous structural characterisation (**Supplementary** Fig. 1A ) [39]. Group 1 mAbs, which bind to the interior face of RBD, were able to neutralize when interior epitopes I-1, I-2 and I-3 were reverted to WT. Group 4 mAbs, which are known to bind to the outer face of the RBD, were rescued by revertant mutant O-1.

Neutralization by REGN10987 was not rescued by any of the revertant spikes in the panel (**Supplementary** Fig. 1B ), suggesting a different combination or additional reversions are needed to restore neutralization sensitivity. Sotrovimab, in contrast to the other mAbs tested, could weakly neutralize XBB.1.5+ (IC_50_ of 0.31 µg/mL). Although sotrovimab is known to bind the outer face of RBD (**Supplementary** Fig. 1C )[40], an increase in neutralization was observed with reversion of residues on the interior face I-1, I-2 and I-3. This suggests that the antigenic landscape of RBD and/or accessibility to neutralizing epitopes, may be influenced by mutations distal to the antibody binding site. This was also seen for several Group 3 RBM mAbs that showed neutralizing activity with reversion of residues in the distal I-1 epitope. To determine if these epitope reversions affected neutralization of mAbs binding to epitopes outside of the RBD, several NTD binding mAbs and an SD1 binding mAb[41] were tested against the revertant virus panel. Neutralization by NTD nAbs was uneffected by reversions in the RBD (**Supplementary** Fig. 2 ). P008_60, an SD1 binding neutralizing antibody with modest cross-neutralization of SARS-CoV-2 variants[41] showed slightly improved neutralization when mutations on the interior of the RBD were reverted, suggesting that despite their distance from SD1, mutations occurring in XBB1.5 may impede accessibility to the SD1 epitope. To investigate whether RBD reversion mutations influence ACE2 binding and therefore subsequent sensitivity to antibody-mediated neutralization, we determined IC_50_ values for soluble ACE2 neutralization for the virus revertant panel (**Supplementary** Fig. 3 ). A previous study has shown that higher neutralisation by soluble ACE2 correlated with ACE2 binding affinity[28]. There was minimal variation in IC_50_ values for the majority of the virus revertant panel when compared to D614G and XBB.1.5+, suggesting that similar levels of ACE2 binding between Spikes. I-3 revertant had reduced neutralisation by soluble ACE2 which may be attributed to the reversion of Q498R and N501Y, both of which have been shown to increase ACE2 affinity in some variants[29]. However, all revertant Spikes gave high titres in the pseudotyped virus neutralization assay, suggesting that virus infectivity was not significantly impeded by ACE2 binding affinity.

### Primary infection cohort

Having successfully generated a panel of XBB1.5+ revertant viruses we next used these as a tool to examine the epitope immunodominance following primary infection with different variants of concern (VOCs) in unvaccinated individuals (**Supplementary Table 3** ). It has previously been observed that the variant an individual is first exposed to through infection can impact neutralization breadth of the resultant antibody response[42]. However, differences in epitope immunodominance elicited following different primary infections have not been extensively studied. Using sera from individuals infected in wave 1 or with the Delta or BA.1 variants (determined by viral sequencing), we examined differences in neutralization breadth against VOCs (**Figure 2A**) and the neutralization of the XBB.1.5+ revertant virus panel (**Figure 2B**). When considering neutralization breadth of convalescent sera against VOCs, ID_50_ values were highest for the autologous infection variant (**Figure 2A**). Wave 1, and Delta convalescent sera showed a sharp drop in ID_50_ against BA.1 highlighting the increase in antigenic distance between pre-Omicron variants and Omicron sub-lineages[43]. Equally, BA.1 convalescent sera had a 22- to 49-fold decrease in neutralization potency against pre-Omicron variants.

**Figure 2.**
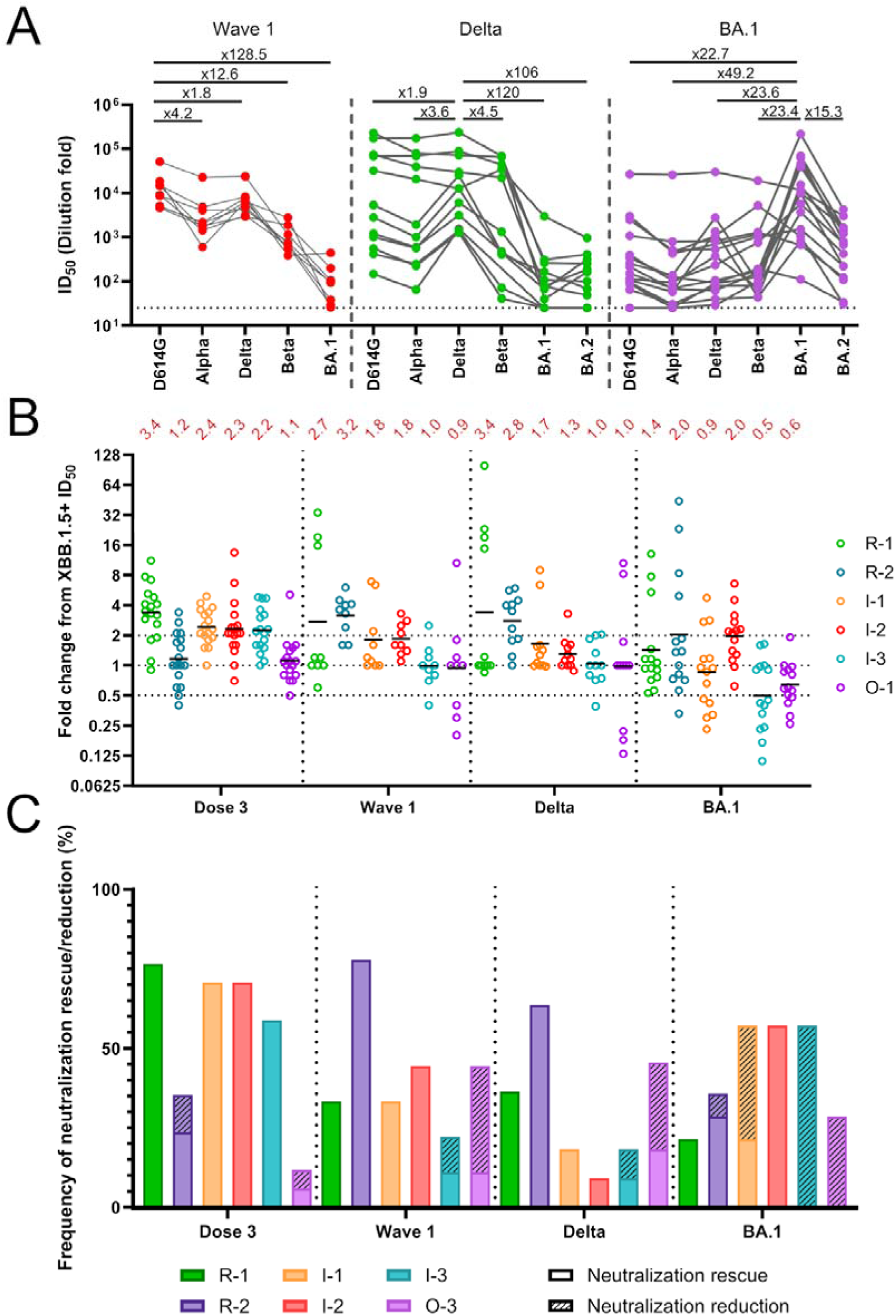
Epitope immunodominance in sera following primary infection with SARS-CoV-2 variants of concern. (A) Cross-neutralizing activity in sera from individuals infected in either wave 1 or with Delta or BA.1. Sera were collected in the acute phase of infection (12-28 days post infection). Each dot represents one individual, and matched data points for neutralization of different VOCs by the same serum sample are connected by lines. Fold reduction in neutralization from the autologous virus is shown above each panel. **(B)** Fold change in ID_50_ of revertant viruses compared to XBB.1.5+ for each individual (Fold change = ID_50_ revertant/ID_50_ XBB.1.5+). Individuals are grouped based on the vaccine status or infecting variant on the X-axis and divided by dotted lines. Neutralization rescue is defined by a fold change in ID_50_ from XBB.1.5+ of greater than 2. Neutralization reduction is defined by a fold change in ID_50_ from XBB.1.5+ less than 0.5. (Dose 3 (*n*=17), wave 1 (*n*=9), Delta (*n*=11), BA.1 (*n*=14); where *n* refers to number of individuals in each group). **(C)** Bar graph showing percentage frequency of donors showing neutralization rescue (solid fill, >2-fold change) and reduction (line pattern, <0.5-fold change).

To quantify and compare epitope usage in the convalescent sera, the fold change in ID_50_ compared to XBB.1.5+ was determined for each revertant virus and these were compared to sera from individuals receiving 3-doses of BNT162b2 mRNA vaccine (**Figure 2B** ). Comparing the geometric means of the fold change showed distinct immunodominance profiles were elicited following vaccination or primary infection by different variants. Of note, BA.1 sera showed higher neutralization of XBB.1.5+ mutant compared to the vaccinee, wave 1 and Delta groups (**Supplementary** Fig. 4 ) which were reflected in lower absolute fold changes for each revertant. This was presumably due to reduced antigenic distance between XBB.1.5+ and BA.1 compared to the pre-omicron variants[43]. A fold change >1 would indicate a neutralizing epitope had been restored within the revertant Spike. This scenario is most relevant for sera from individuals vaccinated with a Wuhan-1 based vaccine or exposed to a variant more similar to Wuhan-1. In contrast, a fold-change <1 would indicate a neutralizing epitope had been removed from a revertant Spike. This scenario is most relevant for sera from individuals exposed to variants that are more similar to XBB.1.5 and was observed in BA.1 primary infection, where responses to I-3 and O-1 gave geometric mean fold changes of 0.5 and 0.6, respectively. To better capture both forms of epitope usage (i.e rescue and reduction), fold change values above 2 and below 0.5 were defined as neutralization rescue and neutralisation reduction, respectively.

The percentage frequency of sera with neutralization rescue or reduction (i.e. >2 or <0.5-fold change) were grouped by epitope reversion for each variant primary infection and compared with 3-dose WT mRNA vaccine sera (**Figure 2C**). Each group showed distinct patterns of epitope usage. Vaccinee sera predominantly targeted the R1 and interior (I-1, I-2, and I-3) epitopes. Although the Spike sequences in wave 1 are almost identical to the vaccine strain, epitope immunodominance was skewed towards R-2 epitope instead of R-1. The reduced R-1 epitope usage in infection sera may suggest a difference in Spike/RBD presentation on the virus surface compared to the vaccine encoded Spike that may be attributed to the stabilising mutations introduced. A skewing towards R-2 and to a lesser extent R-1 and O-1 was observed in Delta convalescent sera with little response against interior epitopes (I-1, I-2 and I-3) suggesting a narrower response. Interestingly, neutralization reduction was seen for O-1 which reverts the mutation at position 452 which is naturally present in Delta Spike and may suggest presence of nAbs dependent on amino acid R452. Primary infection by the more antigenically distant BA.1 elicited an immune response primarily targeting the interior epitopes of RBD and to a lesser extent O-1. However, whilst I-2 showed increased neutralization, I-1, I-3 and O-1 showed decreased neutralization. The I-1, I-3 and O-1 revertants encode mutations at positions 417, 498/501/505, and 339/440, respectively that are naturally present in BA.1 and may therefore also indicate presence of antibodies targeting BA.1 specific residues/epitopes. Overall, the variant responsible for primary infection strongly influences the neutralizing epitopes targeted on Spike.

### Immune imprinting from vaccination influences BTI immunodominance

We next sought to understand how prior vaccination might influence the immune response to variant infection. To study this, neutralization activity against the virus revertant panel was determined for longitudinal serum samples collected from individuals following their 2^nd^ and 3^rd^ doses of mRNA vaccine, as well as 3-4 weeks and 6 months post BTI (**Figure 3A, Supplementary Table 3** )[44]. Results were first assessed by examining the fold change in neutralization potency for each revertant virus compared to XBB.1.5+ over time (**Figure 3A**). The average fold rescue after two vaccine doses was generally low, with R-1, I-1 and I-2 showing the largest rescue (range 1.8-2.2). However, there was an increase in neutralization rescue between the 2^nd^ and 3^rd^ vaccine dose for all revertants, except O-1. The RBM revertant R-1 and all interior RBD revertants (I-1, I-2 and I-3) had neutralization rescue greater than 2-fold (geometric mean). Following BTI, there was increased heterogeneity in the responses between individuals. When considering the average fold change in neutralization for each revertant, the R-1 epitope was the most immunodominant with a geometric mean fold change of 4.5 after BTI. Epitopes R-2 and O-1 had the lowest neutralization rescue, suggesting these regions were not major targets of the neutralizing response after vaccination and BTI.

**Figure 3:**
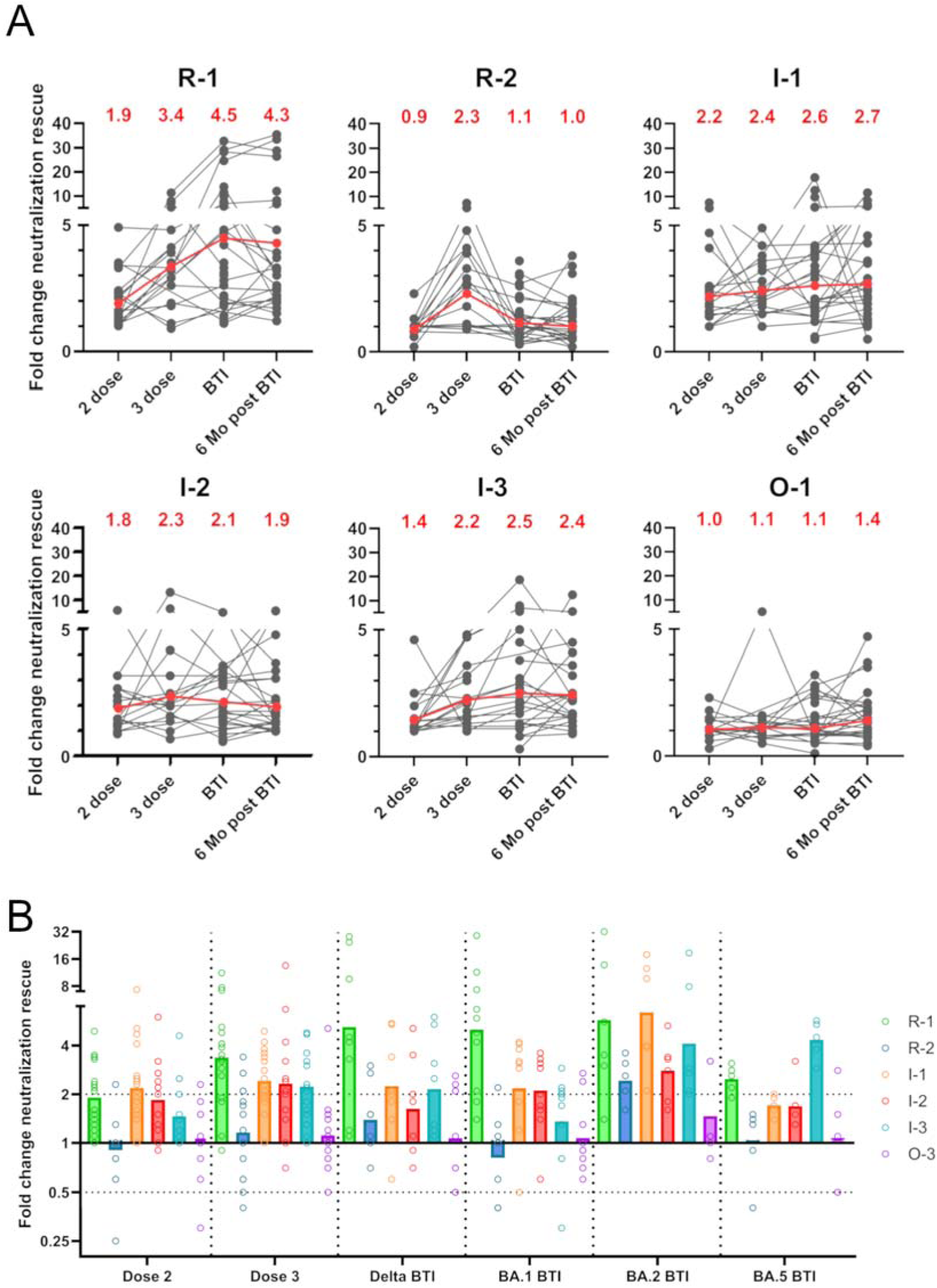
Epitope immunodominance and immune imprinting in sera following mRNA vaccination and BTI. (A) Fold change in ID_50_ of revertant viruses compared to XBB.1.5+ for sera collected following 2- and 3- vaccine doses and 3-4 weeks and 6 months following BTI. Grey lines are donor-matched ID_50_ across four different time points. Geometric mean fold changes are shown at the top of each panel and as red lines. (B) Comparison of the fold change in ID_50_ of revertant viruses compared to XBB.1.5+ for individuals grouped by presumed BTI variant or vaccine status. Bars indicate geometric mean fold change across all individuals in the group. (Dose 2 (^n^=17), Dose 3 (^n^=17), Delta BTI (^n^=8), BA.1 BTI (^n^=10), BA.2 BTI (^n^=6), BA.5 BTI (^n^=5); where ^n^ refers to number of individuals in each group).

### Antigenic distance and time post-exposure may override immune imprinting

As analysis of timepoints post BTI had shown increased heterogeneity, the cohort was regrouped based upon the variant that led to the BTI (determined based on date of positive COVID-19 test). This revealed distinct patterns of epitope usage between infecting variants (**Figure 3B** ). The patterns of neutralization rescue following Delta and BA.1 BTI were similar to that of sera following 2- or 3- doses of vaccine. These sera had R-1 being the most immunodominant epitope and to a lesser extent I-1 and I-2. Sera from individuals with BTI from the more antigenically distant variants, BA.2 and BA.5, showed a shift in immunodominance compared to the vaccine groups (Dose 2 and Dose 3). The BA.2 group showed greatest rescue for R-1, I-1 and I-3 and the BA.5 group showed an immunodominance shift from R-1 towards I-3. These data suggest that a greater antigenic distance of a variant Spike may be better able to override initial immune imprinting from the vaccine. A factor that could also influence immune imprinting is the time between final vaccination and time of BTI[42]. Those presumed to be infected with BA.2 or BA.5 had a longer period between vaccination and BTI compared to Delta and BA.1 BTI groups (**Supplementary Table 3**).

### Primary infection epitope usage differs from BTI

Next, we compared the epitope immunodominance between primary infection and BTI for Delta (**Figure 4A**) and BA.1 (**Figure 4B**). The percentage frequency of individuals showing a >2-fold increase (neutralization rescue) or <0.5-fold decrease (neutralization reduction) in ID_50_ were plotted. Immunodominance in Delta and BA.1 BTI were similar to the response after 3 vaccine doses (**Figure 3C**) and were largely skewed towards R-1, I-1, I-2 and I-3 epitopes.

**Figure 4:**
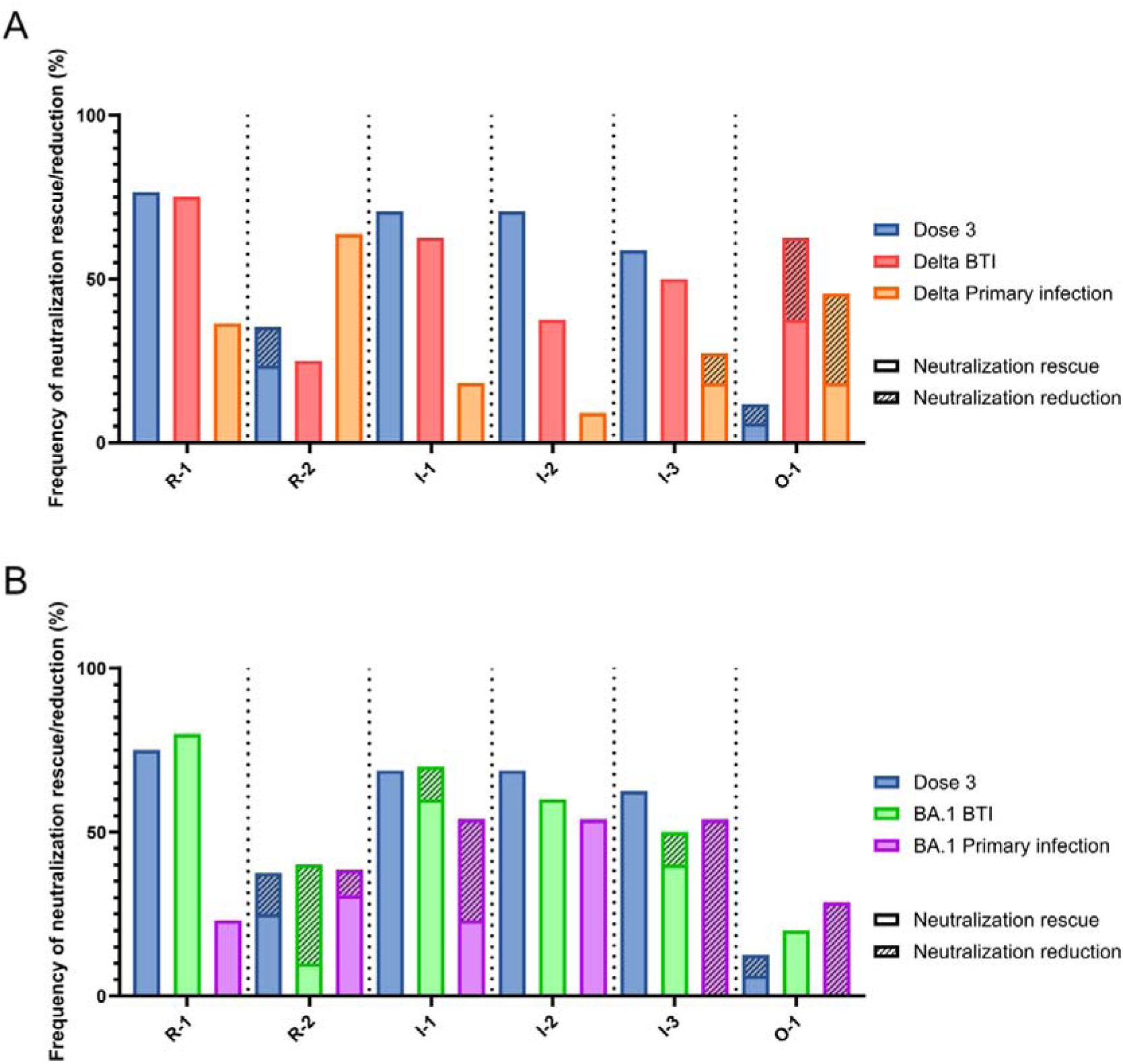
Comparison of immunodominance between primary infection and BTI. Bar graph comparing percentage frequency of donors showing neutralization rescue (solid fill, >2-fold change) and reduction (line pattern, <0.5-fold change) for sera following 3 vaccine doses, BTI or primary infection. (A) Shows data for Delta BTI and Delta primary infection. (Dose 3 (^n^=17), Delta BTI (^n^=8), Delta primary infection (^n^=11); where ^n^ refers to number of individuals in each group). (B) Shows data for BA.1 BTI and BA.1 primary infection. Comparisons are grouped by reverted epitope on the X-axis and separated by dotted lines. (Dose 3 (*n*=17), BA.1 BTI (*n*=14), BA.1 primary infection (*n*=10); where *n* refers to number of individuals in each group).

Interestingly, the pattern of immunodominance in Delta and BA.1 primary infection sera differed from that in BTI. For Delta infection, there was a decrease in frequency of individuals with neutralization rescue in epitope R-1, I-1 and I-2 and there was a shift towards the R-2 epitope. However, both Delta BTI and primary Delta infection both showed individuals with a reduced neutralization to the O-1 revertant. This suggests the O-1 epitope may be a target of variant specific antibodies, elicited by activation of naïve B-cells by Delta variant infection. Mutation at position 452 is present in the Delta variant (L452R) and the O-1 revertant and has been shown to be highly immune evasive [45]. This may suggest that a *de novo* response has been developed towards R452 following BTI, overriding the humoral immune imprinting from the vaccine.

For BA.1 infection, there was also a decrease in frequency of individuals with rescue of R-1 compared to BA.1 BTI. Interestingly, for I-1, I-3 and O-3 there were contrasting neutralization capabilities between BTI and primary infection sera. Whilst serum neutralization was rescued in these revertants following BA.1 BTI, the response elicited by BA.1 primary infection was predominantly attributed to neutralization reduction. This suggests that primary infection elicits a more BA.1-specific neutralizing response which have been supressed in the BA.1 BTI group.

## Discussion

Our resistant variant and virus revertant panel approach provides an accessible and potentially high-throughput method to map the epitopes targeted by neutralizing antibodies in sera. Furthermore, this approach can be quickly adapted to newly emerging variants of SARS-CoV-2. The ability to determine epitope immunodominance without the need to have access to PBMCs for mAb isolation allows for wider application across larger cohorts. Additionally, we can directly interpret the functional effects of Spike mutations by measuring neutralizing activity of sera as opposed to only binding. In the context of polyclonal serum this is an advantage as a notable proportion of antibodies in sera can bind but lack neutralizing activity[2,46].

Using our revertant virus panel we have mapped the immunodominant sites of RBD in sera following primary infection and across longitudinal time points following vaccination and BTI. When considering vaccinee sera alone, the mutations in XBB.1.5+ that facilitated neutralization rescue were found in 5 out of 6 revertant epitopes, including within both the RBM (R-1 and R-2) and inner face of RBD (I-1, I-2 and I-3). This shows the polyclonal nature of the antibody response to SARS-CoV-2 vaccination and the selective pressure these nAbs have on emerging of variants. By focusing on mutations derived from those naturally occurring in VOCs we can link immunodominance at a given time point to viral escape. Indeed, mutations in the R-1 revertant, that correspond to those that arose in Omicron and its sub-lineage variants, may have been driven by the high frequency of serum antibodies targeting the RBM following vaccination.

When the immunodominance profile for vaccine-induced antibody responses was compared to that following primary infection with different variants, distinct epitope usage was observed. Despite the BNT162b2 vaccine and wave 1 infections presenting almost identical Spikes, a narrower response primarily targeting R-2 was observed following primary infection. There was also a reduced response towards epitopes on the interior face (I-1, I-2, and I-3) compared to vaccine-induced antibodies. Similar results have previously been reported by Greaney *et al,* in which DMS was utilised to compare specificity of serum antibodies from mRNA-1273 vaccination and SARS-CoV-2 infection[31]. Vaccine-induced antibodies were shown to target a broader range of epitopes on the RBD when compared to infection-induced serum antibodies. These results suggest that presentation and conformation of Spike expressed by mRNA vaccination may differ from Spike in its native conformation on the viral membrane. The interior face of the RBD is immunodominant in vaccinee sera, however, these epitopes are only accessible when RBD is in its “up”, conformation[47]. The addition of stabilising mutations (K986P and V987P) to mRNA vaccine Spike sequences, have been speculated to influence RBD conformation[48]. Whether these mutations influence immunodominance by modulating RBD conformation requires further investigation. Indeed, these stabilising mutations were not incorporated in the ChAdOx1 nCoV-19 vaccine and it is possible that the use of different vaccine platforms may also influence epitope immunodominance to RBD[49].

Several groups have shown, using B cell staining, that variant infection/boosting in vaccinated individuals preferentially re-engages pre-existing memory B cells and suppresses generation of *de novo* variant-specific responses[14,22,50]. Similarly, we show that immune imprinting from mRNA vaccines is persistent following multiple vaccine doses and BTI[14,51,52]. However, by further characterising imprinted responses, we reveal more subtle shifts in immunodominance following BTI. Antigenic distance and the waning antibody response over time seem to be factors in the persistence of immune imprinting. Indeed, BTI with later omicron sub-lineages (BA.2 and BA.5 variants) shifted immunodominance towards epitopes on the interior face of RBD. Similar to the vaccine strains, this may be explained by mutations in the omicron lineage stabilizing the spike in its RBD “up” conformation[53], hence increasing accessibility of the interior face to antibody binding.

Distinct epitope usage was also observed in convalescent sera from individuals infected with Delta and BA.1 variants, highlighting their antigenic distance from the Wuhan-1 strain. When compared to BTI with the matching variant, the influence of imprinting from the vaccine was apparent. Epitope immunodominance following BTI predominantly favoured the imprinted response elicited by the vaccine. However, some evidence of antibody responses to variant-specific epitopes was observed. The L452R mutation, present in the Delta variant seems to elicit a *de novo* antibody response able to override imprinting. Additionally, BA.1 mutations on the interior face of RBD were able to elicit a variant-specific response. These findings were supported at a monoclonal level, where individuals with Delta and BA.1 BTI had a higher frequency of antibodies targeting the group 4 epitope (located on the outer face of RBD and represented by mutations in the O-1 revertant) than individuals with primary infection during wave 1 (**Supplementary** Figure 5 )[2,23]. A study by Guerra *et al* isolated a panel of monoclonal antibodies from a P.1 infected individual[54]. Interestingly, they identified a mAb, COVA309-03, that was specific for the 484K residue present in P.1 which supports our observation that there are nAbs targeting variant specific mutations in the sera of primary Delta and BA.1 infected individuals.

Although vaccines can be rapidly adapted to new variants that arise, a large proportion of the population was vaccinated with the Wuhan-1 SARS-CoV-2 Spike. Conversely, looking towards the future, new generations of naïve infants will likely not be exposed to the ancestral variant or past variants of SARS-CoV-2. This new generation will be imprinted with variant vaccines or the current circulating VOCs and therefore likely have a differently targeted neutralizing antibody response compared to those studied so far. The heterogeneity of exposure histories in the general population and the effects on immune imprinting by prime/boosting with new variant-based vaccines will need to be considered in the design of updated vaccines to ensure efficient protection.

## Methods

### Ethics statement

This study used human samples collected with written consent as part of a study entitled “Antibody responses following COVID-19 vaccination.” Ethical approval was obtained from the King’s College London Infectious Diseases Biobank (IBD) (KDJF-110121) under the terms of the IDB’s ethics permission (REC reference: 19/SC/0232) granted by the South Central Hampshire B Research Ethics Committee in 2019 and London Bridge Research Ethics Committee (reference: REC14/LO/1699). Collection of surplus serum samples at St Thomas Hospital, London, was approved by South Central-Hampshire B REC (20/SC/0310).

### Description of human samples

#### Primary infection cohort

Surplus serum samples from 37 individuals with primary infection were collected at St. Thomas’ Hospital London. SARS-CoV-2 cases were diagnosed by RT-PCR of respiratory samples and infecting variants were confirmed using whole genome sequencing as previously described[42], or using MT-PCR[55]. The cohort were 54% male, 41% female and 5% undetermined with a median age of 57 (range 7-90 years) (**Supplementary Table 3** ). Samples were collected at a median of 18 days (range 12-28 days) post onset of symptoms. 27% (9/34) individuals were infected during wave 1 (presumably with variants close to the ancestral Wuhan-1 strain), 32% (11/34) were infected with Delta variant and 41% (14/34) were infected with BA.1 variant.

#### Vaccine and BTI cohort

Sequential serum samples were collected from a cohort of 44 donors, spanning 3 doses of BNT162b2 mRNA vaccine and BTI with approval granted by South Central – Hampshire B (REC reference: 19/SC/0232). The cohort were 45% male and 55% female. SARS-CoV-2 infection was diagnosed by RT-PCR and/or lateral flow testing and confirmed by presence of IgG to nucleoprotein. Infecting variant was estimated by the dominant strain circulating in the UK at the time of infection. 18% (8/44) individuals were infected during the Delta wave, 32% (14/44), 18% (8/44) and 14% (6/44) were infected during BA.1, BA.2 and BA.5 waves, respectively. 7% (3/44) donors were infected during periods of transition between dominant variants, hence infecting variant was undetermined. 11% (5/44) individuals recorded no BTI. Time post 3rd vaccine dose varied for each group with most of the Delta breakthrough infection occurring prior to the 3rd vaccine dose (median -59 days). BA.1, BA.2, and BA.5 BTI occurred at a median of 84-, 136-, and 246- days post 3rd dose, respectively.

### Spike mutagenesis

Spike mutants were generated with Q5 Site-Directed Mutagenesis Kit (NEB, E0554) following the manufacturer’s instructions. XBB.1.5+ was generated by introducing L452R (forward primer TACAATTACCGGTACCGGCTGTTC, reverse primer GTTGCCGCTGGGTTTGGA) and K444T (forward primer CTGGACTCCACACCCAGCGGC, reverse primer CTTGTTGCTGTTCCAGGCAATC) mutations into the XBB.1.5 spike sequence. Primer sequences for reversion mutations are shown in (**Supplementary Table 1** ). Sanger sequencing was used to confirm presence of desired mutations.

### Pseudovirus production

Pseudotyped HIV-1 virus incorporating each of the mutated SARS-CoV-2 Spikes were generated by first seeding HEK293T/17 cells in 10 cm dishes at a density of 7x10^5^ cells/mL in Dulbecco’s Modified Eagle Medium (DMEM) with 10% fetal bovine serum (FBS) and 1% Pen/Strep. Following overnight culture, cells were co-transfected using 90 µg PEI-Max (1 mg/mL, Polysciences) with 15 µg HIV-luciferase plasmid, 10 µg HIV 8.91 gag/pol plasmid, and 5 µg SARS-CoV-2 Spike protein plasmid[56]. Transfected cells were incubated for 72 h at 37°C, and virus was harvested, sterile filtered, and stored at −80°C until required. Mutations present in each variant Spike are shown in (**Supplementary Table 2**).

### Human ACE2 expression

Recombinant human ACE2 containing a Twin-StrepTag was expressed in Expi293F cells (Invitrogen). 1000 µg of plasmid DNA was transfected with PEI-Max (1 mg/mL, Polysciences), at a 1:3 ratio, in 1 L of cells at 1.5x10^6^ cells/mL density. Supernatant was harvested and protein was purified with SrepTactinXT Superflow high-capacity 50% suspension (IBA Life Sciences) according to the manufacturer’s protocol by gravity flow.

### Pseudovirus neutralisation assays

In half-well 96 well plates, sera, mAb or soluble ACE2 was serially diluted in DMEM (10% FBS, 1% Pen/Strep) with a starting serum dilution of 1:10 dilution or mAb concentration 25 µg/mL. Pseudotyped D614G or XBB.1.5+ variant viruses was added and incubated for 1 hr at 37°C. Hela- ACE2 cells were added at a density of 4x10^5^ cells/mL and incubated for 72 hr at 37°C. Levels of infection were measured by luminescence with Bright-Glo luciferase kit (Promega) on a Victor X3 multilabel reader (Perkin Elmer). Samples were run in duplicate to determine IC_50_ and ID_50_ values. Fold change in neutralization against reverted mutants was calculated as a fold change of IC_50_ or ID_50_ from XBB.1.5+. Statistical analyses were performed using GraphPad Prism v.10.2.1.

## Acknowledgements

We thank Wendy Barclay for providing the Beta, Delta, BA.1, BA.2, and XBB.1.5 Spike plasmids, Peter Cherepanov for providing the ACE2 expression plasmid, and James Voss for providing the HeLa-ACE2 cells.

