## Supplementary for "Profiling serum immunodominance following SARS-CoV-2 primary and breakthrough infection reveals distinct variant-specific epitope usage and immune imprinting"

**Supplementary Table 1:** Primer sequences for XBB.1.5+ Spike mutagenesis

| **Name** | **Forward primer** | **Reverse primer** |
| --- | --- | --- |
| **L452R** | TACAATTACCgGTACCGGCTGTTC | GTTGCCGCTGGGTTTGGA |
| **K444T** | CTGGACTCCAcACCCAGCGGC | CTTGTTGCTGTTCCAGGCAATC |
| **S477T478** | TCAGGCCGGCtcCAcGCCTTGTAACG | TAGATCTCGGTGGAGATG |
| **F486** | GTGGCAGGCTtCAACTGCTAC | GCCGTTACAAGGCGTGGAG |
| **E484** | AACGGCGTGGaAGGCTCCAAC | ACAAGGCTTGTTGCCGGC |
| **F490** | AACTGCTACTtCCCACTGCAGTCC | GGAGCCTTCCACGCCGTT |
| **S375T376** | CGCCCCCTTCtccaCCTTCAAGTG | AAGTTGTAGATCACGGAGTAG |
| **R408** | AAATGAAGTGcGTCAGATTGCCCCTG | CCCCGGATCACGAAGCTG |
| **D405** | GATCCGGGGAgATGAAGTGAGTC | ACGAAGCTGTCGGCGTAC |
| **K417** | AGACAGGCAAgATCGCCGACT | GTCCAGGGGCAATCTGAC |
| **R346** | AATGCCACCAgATTCGCCTCT | GAACACCTCGTGGAAGGG |
| **G339** | GTGCCCCTTCggCGAGGTGTTC | AGATTGGTGATATTGGGG |
| **Q498** | TACGGCTTTCaGCCCACATATG | GGACTGCAGTGGGGAGTA |
| **N501** | TCAGCCCACAaATGGCGTGG | AAGCCGTAGGACTGCAGTGG |
| **Y505** | TGGCGTGGGCtATCAGCCCTA | TTTGTGGGCTGAAAGCCG |
| **N440** | ACAGCAACAAtCTGGACTCCAAAC | TCCAGGCAATCACACAGC |

**Supplementary Table 2:** Mutations present in each reverted XBB1.5+ Spike variant

| **Name** | **Mutations** |
| --- | --- |
| XBB.1.5+ | K444T, L452R (additional mutations) |
| R-1 | N477S, K478T, S486F |
| R-2 | A484E, S490F |
| I-1 | N405D, N417K |
| I-2 | F375S, A376T, S408R |
| I-3 | R498Q, Y501N, H505Y |
| O-1 | H339G, T346R, K440N, K444, L452 |

**Supplementary Table 3:** Cohort descriptions for primary infection cohort and Longitudinal vaccine/infection cohort. N/A: not applicable.

|  | **Primary infection cohort n=34** | **Longitudinal cohort n=44** |
| --- | --- | --- |
| **Age (years)** | 7-90 (median = 58) | 22-63 (median = 33) |
| **Gender** |  |  |
| Male | 59% (20/34) | 45% (20/44) |
| Female | 35% (12/34) | 55% (24/44) |
| Undetermined | 6% (2/34) | 0% (0/44) |
| **Primary infection** |  |  |
| Wave 1 | 27% (9/34) | N/A |
| Delta | 32% (11/34) | N/A |
| BA.1 | 41% (14/34) | N/A |
| **Days post onset of symptoms  of collection (days)** | 12-28 (median = 18) | N/A |
| **Variant BTI** |  |  |
| Delta | N/A | 18% (8/44) |
| BA.1 | N/A | 32% (14/44) |
| BA.2 | N/A | 18% (8/44) |
| BA.5 | N/A | 14% (6/44) |
| Undetermined | N/A | 7% (3/44) |
| No BTI | N/A | 11% (5/44) |
| **Time of BTI post 3rd dose (days)** |  |  |
| Delta | N/A | -59 |
| BA.1 | N/A | 84 |
| BA.2 | N/A | 136 |
| BA.5 | N/A | 246 |

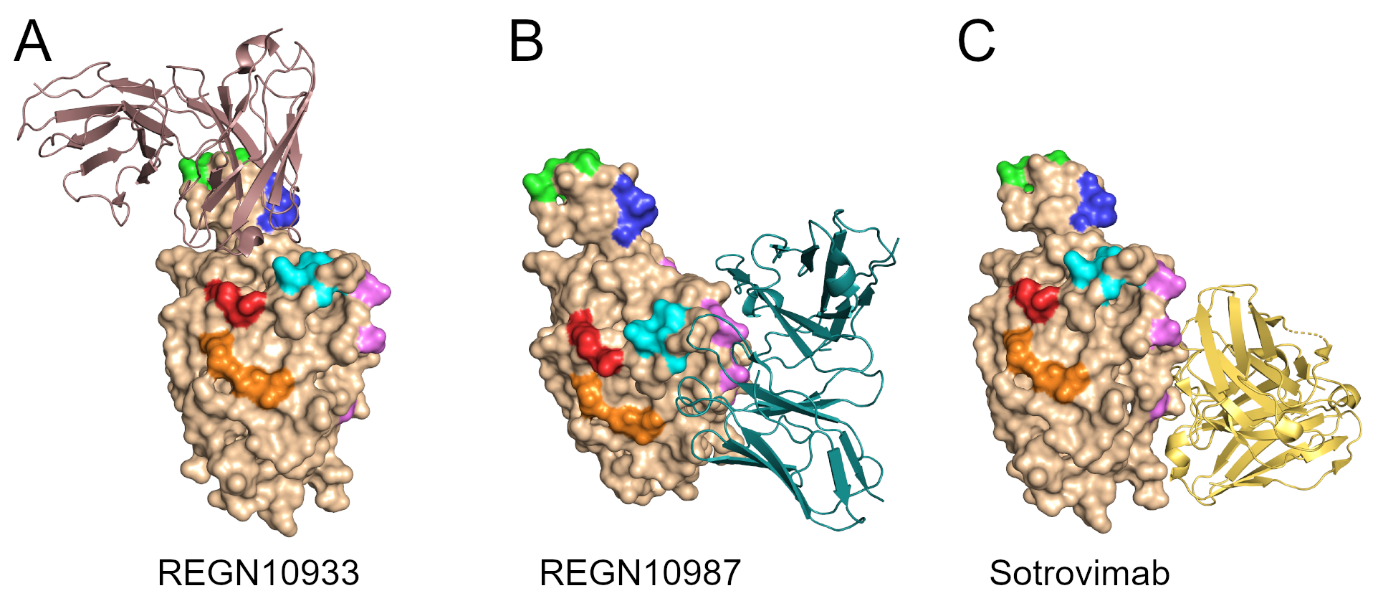
**Supplementary Figure 1**: Structures of therapeutic monoclonal antibodies in complex with RBD; **(A)** REGN10933 (PDB: 6XDG), **(B)** REGN10987 (PDB: 6XDG) and **(C)** Sotrovimab (PDB: 6WPS). Fabs (cartoon representation) are binding to SARS-CoV-2 RBD (surface representation) with revertant residues coloured as in **Figure 1C** and **Supplementary Table S2**.

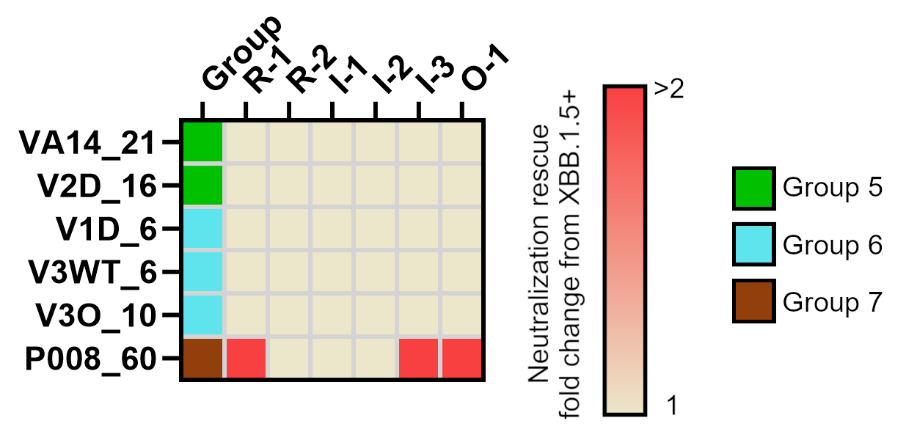
**Supplementary Figure 2**: Heatmap showing fold rescue of neutralisation for reverted viruses compared to XBB.1.5+ for a panel of NTD-specific mAbs (Group 5 and 6) and SD1-specific mAb (Group 7) [1–3].

**Supplementary Figure 3**: Neutralization of pseudoviruses with variant and epitope reverted Spikes by soluble monomeric ACE2. Spikes with higher binding affinity are neutralized more efficiently by soluble ACE2[4]. A table of ACE2 IC_50_ values for each spike is included in the legend.

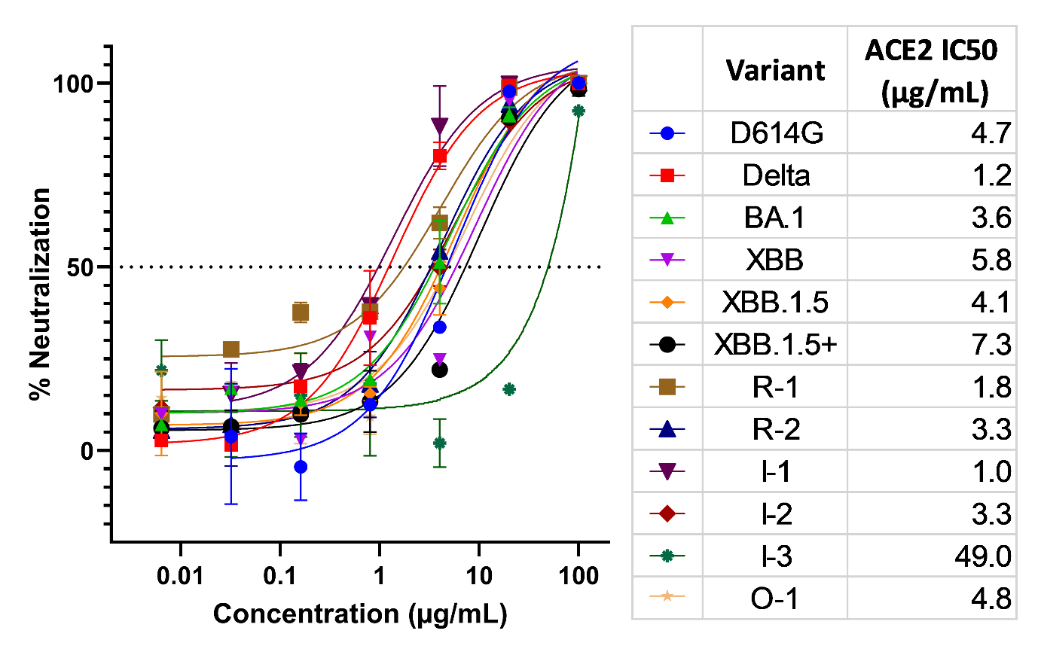

**Supplementary Figure 4:** ID_50_ of sera from longitudinal vaccine and primary infection cohorts against the XBB.1.5+ neutralization resistant mutant pseudovirus (Kruskal-Wallis multiple comparison test, *P < 0.0332, **P < 0.0021, ***P < 0.0002, and ****P < 0.0001). Geometric mean ID­_50_ are indicated by horizontal bars and the fold change compared to BA.1 primary infection sera is shown in red text.

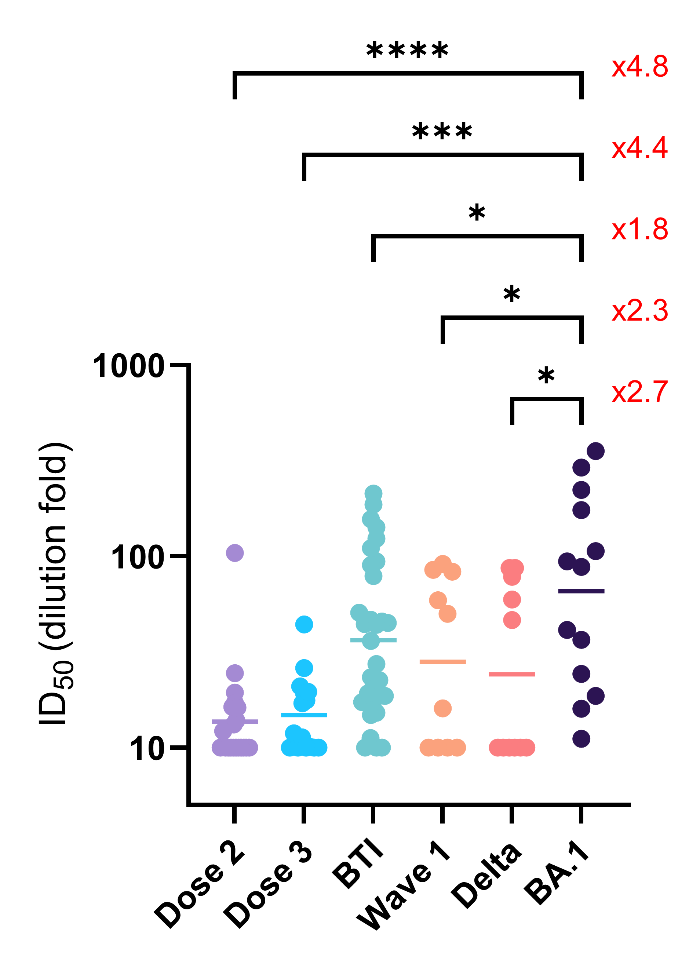

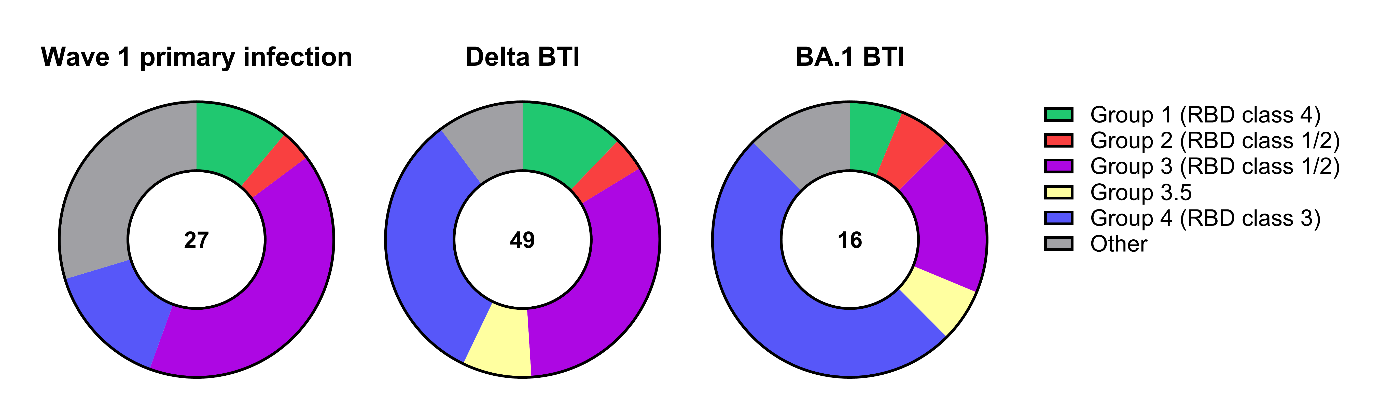
**Supplementary Figure 5:** Pie charts showing epitope usage by mAbs isolated from individuals with primary infection during wave 1[5] and BTI with either Delta and BA.1 variants[2]. Total number of mAbs analysed is shown within the pie chart. RBD groups have been defined previously in Graham *et al*.[5] and matching RBD epitope classes have defined by Barnes *et al.*[6]
